# HP1 binding creates a local barrier against transcription activation and persists during chromatin decondensation

**DOI:** 10.1101/2024.12.09.627308

**Authors:** Robin Weinmann, Arjun Udupa, Jan Fabio Nickels, Lukas Frank, Caroline Knotz, Karsten Rippe

## Abstract

Mouse pericentric repeats form transcriptionally silent and compacted domains known as chromocenters. These prototypic heterochromatin compartments are marked by heterochromatin protein 1 (HP1). However, its contributions to chromocenter structure and function remain debated. We investigated the role of HP1α by recruiting the activators VP16, p65, and VPR to mouse fibroblast chromocenters and analyzed its silencing activity with a transcription reporter. Upon chromocenter decondensation and transcription activation, interactions of HP1α with chromatin and H3K9 trimethylation remained stable, suggesting stoichiometric binding rather than higher-order assembly. HP1α-mediated repression required promoter-proximal binding and effectively suppressed VP16-triggered activation but not the stronger activation by VPR. These observations are explained by a 1D lattice binding model, which conceptualizes chromocenters as arrays of repeat units that can independently switch between silenced and activated states. Our findings provide a quantitative framework that explains how chromocenter organization responds to transcriptional activation while maintaining local heterochromatin features.

**GRAPHICAL ABSTRACT:** 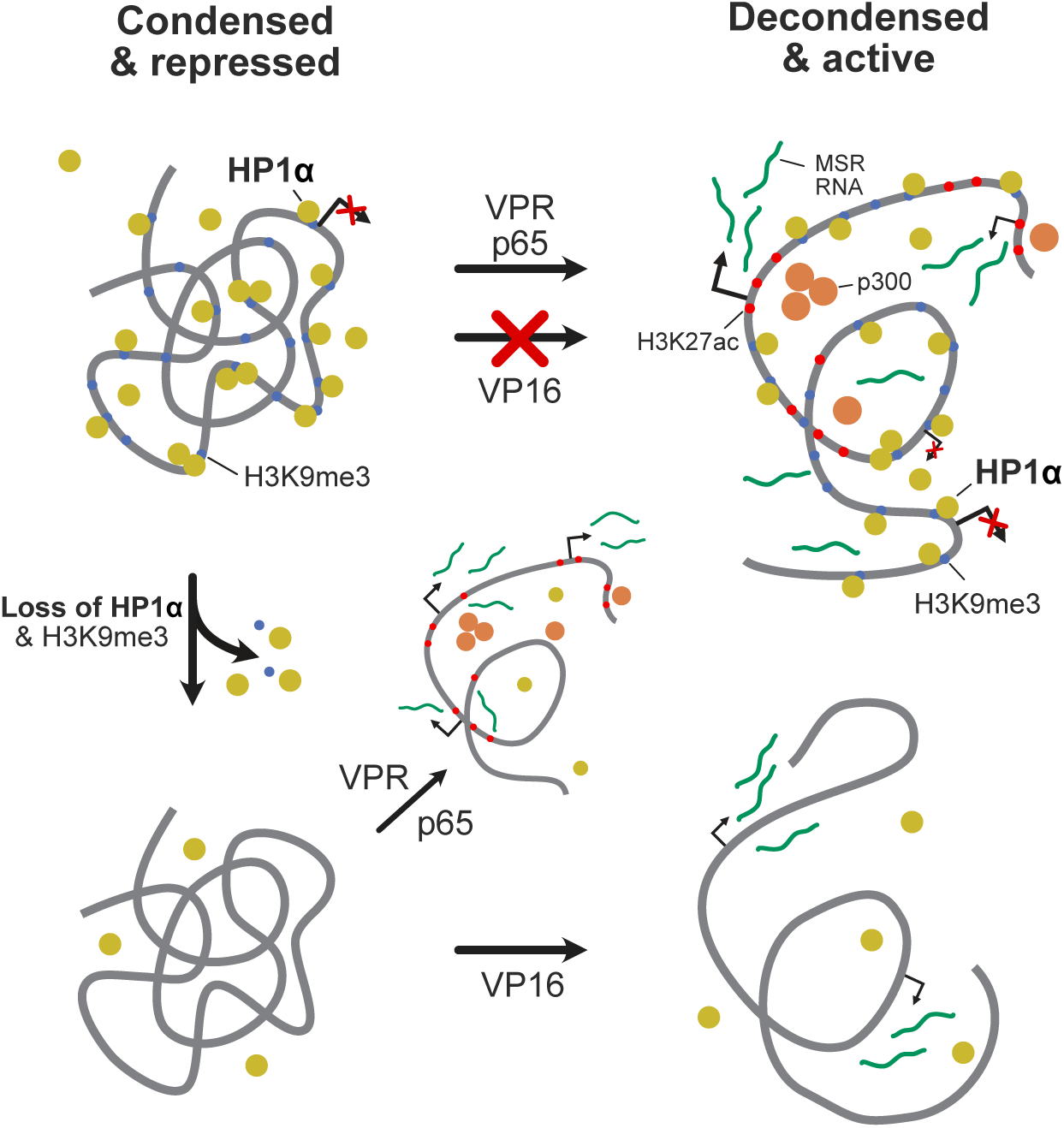

**HIGHLIGHTS:**

- HP1α represses transcription at mouse chromocenters and an ectopic reporter when competing with transcriptional activators VP16, p65 and VPR
- HP1α repression requires promoter-proximal binding and effectively counteracts the weak activator VP16
- The strong activator VPR overcomes HP1α-mediated repression while HP1α remains bound to chromatin
- HP1α binding and H3K9me3 persist during chromocenter decondensation and transcriptional activation
- A 1D lattice binding model explains how independent repeat units transition between silenced and activated states without requiring phase separation

## INTRODUCTION

Heterochromatin protein 1 (HP1) is a crucial marker protein of the compacted and silenced heterochromatin state (Eissenberg and Elgin, 2014; Grewal, 2023; Kumar and Kono, 2020; Kwon and Workman, 2008; Maison and Almouzni, 2004). In human and mouse cells, HP1 exists in three closely related isoforms: HP1α, HP1β and HP1γ. These proteins recognize and bind to tri-methylated histone H3 at lysine 9 (H3K9me3) through their N-terminal chromodomain (CD). Furthermore, HP1 can interact with other proteins like the histone methyltransferases SUV39H1/SUV39H2 (setting H3K9me3) via the PxVxL motif in its C-terminal chromoshadow (CSD) domain. A model system for the interplay of HP1, H3K9me3 and SUV39H1/2 is mouse pericentric heterochromatin, which assembles from thousands of copies of the 234 bp long major satellite repeat (MSR) sequence (Packiaraj and Thakur, 2023; Vissel and Choo, 1989). The resulting transcriptionally inactive chromatin domains on the 1 µm scale are enriched in HP1, SUV39H1/2, and H3K9me3 and are termed chromocenters due to their intense staining with fluorescent dyes like DAPI (Guenatri et al., 2004; Muller-Ott et al., 2014; Padeken et al., 2022; Peters et al., 2001; Probst and Almouzni, 2008). Different models have been proposed that rationalize the formation and maintenance of chromocenters based on binding, feedback loops, and long-range interactions ((Muller-Ott et al., 2014; Thorn et al., 2022) and references therein). One pertinent question is if traditional ligand binding formalisms or polymer models, as used in our previous work (Erdel et al., 2020; Muller-Ott et al., 2014; Thorn et al., 2022), are sufficient to describe HP1 activity in the cell or whether an HP1-driven (liquid-liquid) phase separation (LLPS) is relevant for heterochromatin function (Keenen et al., 2021; Larson et al., 2017; Strom et al., 2017; Tortora et al., 2023). Furthermore, the simple model that HP1 would bind to H3K9me3 and thereby induce chromatin compaction and transcription silencing in an endogenous cellular environment is challenged by several findings: (i) In *Suv39h1*/*Suv39h2* double null mutant (*Suv39h* dn) mouse fibroblasts the enrichment of HP1 and H3K9me3 is lost, but the higher-order organization of the DAPI stained chromocenter structure is unaffected (Erdel et al., 2020; Peters et al., 2001; Schotta et al., 2004). (ii) The knockout of HP1α/β/γ isoforms in mouse lacks structure phenotypes on the 0.1-1 µm mesoscale (Bosch-Presegue et al., 2017; Mattout et al., 2015; Singh and Newman, 2022). (iii) HP1 proteins are present in mouse but absent in adult chicken, frog and fish nucleated erythrocytes, pointing to HP1-independent pathways for heterochromatin formation (Gilbert et al., 2003). (iv) HP1 can act as both an activator and a repressor of transcription suggesting that its activity is context-dependent (Kwon and Workman, 2011; Schoelz and Riddle, 2022).

Despite the prominent role of HP1 as a central heterochromatin marker protein, it is unclear if it compacts chromatin in the cell and how its chromatin binding is related to repeat silencing. It is noted that differences between organisms (e.g., mouse/Drosophila) or developmental stages (early embryo/embryonic stem cells/progenitor/fully differentiated cells) could govern HP1 binding and function. Indeed, changes in HP1 chromatin interaction dynamics and chromatin compaction have been reported in several instances. These include the differentiation of mouse embryonic stem cells (Meshorer et al., 2006), glioblastoma tumor-initiating/differentiated cells (Mallm et al., 2020), local density changes to a more homogeneous state at early differentiation (Joron et al., 2023), and dynamic HP1-dependent heterochromatin states with reduced compaction in pluripotent cells (Dupont et al., 2023).

These observations raise fundamental questions about the role of HP1α in chromocenter organization and function. Here, we dissected the transition of chromocenters from a silenced and compacted to an activated and decondensed state in immortalized mouse embryonic fibroblast (iMEF) cells to address the question of how repeat silencing is linked to HP1α binding and chromatin compaction. This process was studied by recruiting either the transcription activator VP16 from herpes simplex virus (Sadowski et al., 1988), the activation domain of p65 (Schmitz and Baeuerle, 1991) or VPR (VP64-p65-Rta), a tripartite synthetic construct that consists of VP64 (4 copies of VP16) fused to the activation domains of p65 and Rta (Chavez et al., 2015), to the MSR sequences. We elucidated the function and interplay of chromatin compaction/decondensation, HP1α binding, the H3K9me3 and H3K27ac modifications, and MSR transcription. Our results demonstrate that promoter-proximal HP1α binding has a localized repressive activity that potent activators can overcome. The data can be explained by a simple 1D lattice model representing the MSR units and stoichiometric binding of HP1α without invoking a phase separation mechanism.

## RESULTS

We employed two cellular systems to elucidate how HP1α contributes to a repressive heterochromatin environment. Pericentric heterochromatin in chromocenters was studied in wild-type and *Suv39h* dn mouse fibroblast cell lines (**Fig. 1A**). To analyze the gene regulatory activity of HP1α, a reporter array in the human U2OS cell line was used (Janicki et al., 2004). In this system, activators and repressors were recruited in a spatiotemporally defined manner while visualizing and quantifying transcriptional output in living cells (**Fig. 1A**) (Erdel et al., 2020; Rademacher et al., 2017; Trojanowski et al., 2022).

**Figure 1.**
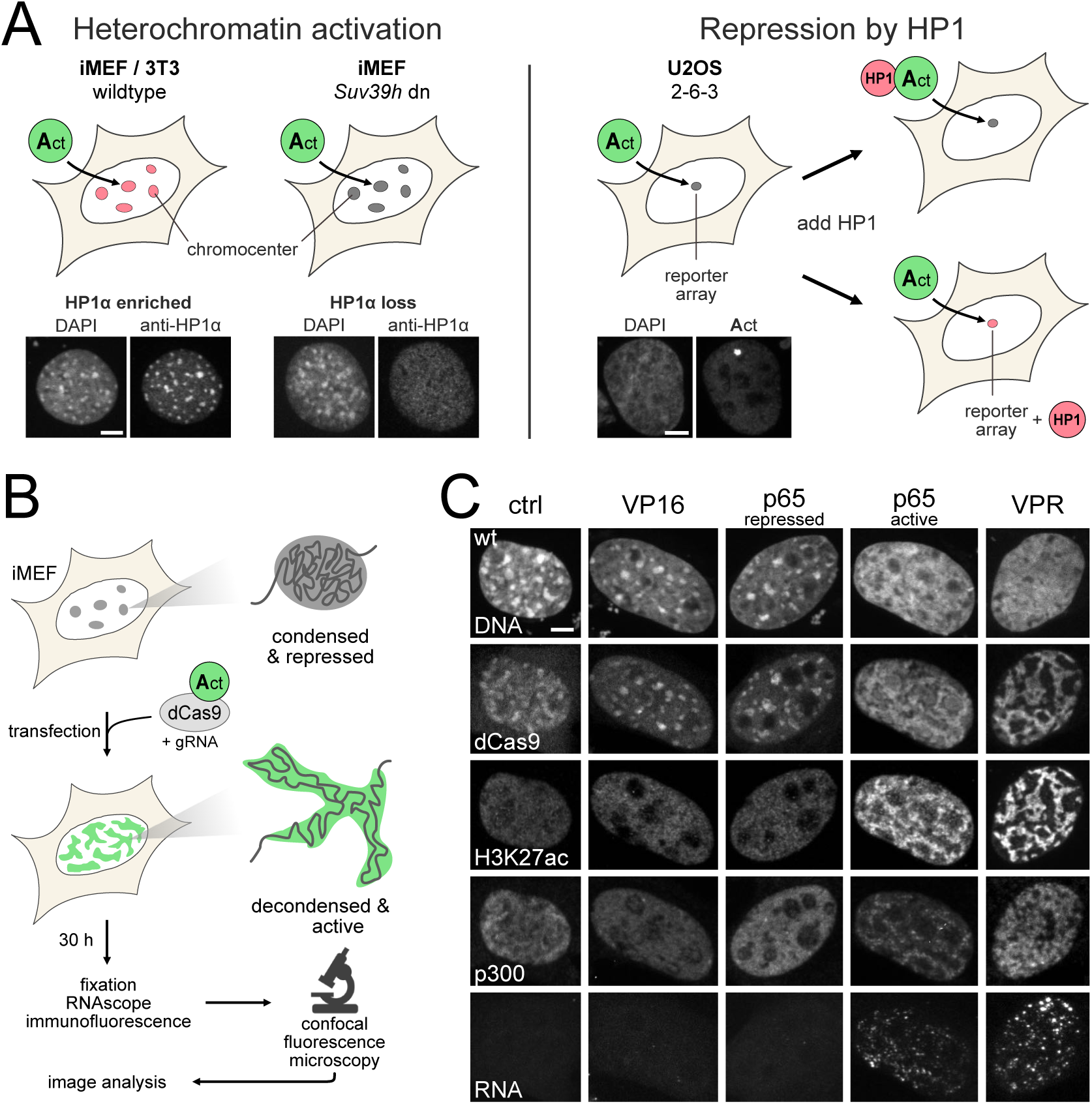
Approaches to assess the repressive function of HP1α. (**A**) Scheme of cellular systems studied. Left: constitutive heterochromatin regions (chromocenters) in iMEFs are targeted by activators to decondense their structure and induce transcription of MSR. Endogenous HP1α is enriched at chromocenters in wild-type cells but not in a *Suv39h dn* cell line, although chromatin compaction persists, as seen in the DAPI-dense foci. Right: activators are recruited together with HP1α in a complex to *tet*O sites or separately to *tet*O and *lac*O sites of the reporter array integrated into the human U2OS 2-6-3 osteosarcoma cell line. The impact of HP1α on transcription induction by the activator is assessed by comparing the transcriptional output between different conditions. Scale bars, 5 µm. (**B**) Scheme of chromocenter activation experiments. Fusion constructs of dCas9-GFP and activator are recruited to chromocenters using sgRNA targeting MSRs. Cells are fixed after 30 h, stained, and imaged using confocal fluorescence microscopy. Images were processed by an analysis pipeline (see **Supplementary Fig. S1**). (**C**) Exemplary confocal fluorescent microscopy images of different activator fusion constructs and their effect on chromocenters. Scale bar, 5 µm.

### VPR decondenses chromocenters and activates transcription of MSR

We combined the methodology introduced in our previous work (Erdel et al., 2020; Frank et al., 2021) to recruit different transcription activators to chromocenters in iMEF cells according to the workflow depicted in **Fig. 1B**. Cells were transfected with plasmids expressing a sgRNA targeting MSRs and plasmids encoding dCas9-GFP fused to one of the three activators VP16, p65 or VPR. Cells were then fixed 30 h after transfection and stained for H3K27ac, p300, and MSR RNA. After visualization by confocal fluorescent microscopy, chromocenters targeted by VPR disassembled from their original spherical structure into dispersed substructures, as reported previously (**Fig. 1C**) (Erdel et al., 2020; Frank et al., 2021). The RNAscope FISH analysis revealed that MSR RNA was produced, indicating that chromocenters in this state no longer silence MSRs. We also observed a high enrichment of the H3K27ac signal colocalizing with dCas9 at MSR sequences and recruitment of the histone acetyltransferase p300 to their vicinity. The co-recruitment of p300 and subsequent H3K27ac deposition could be mediated by the p65 and Rta parts of VPR, which have been shown to interact directly with p300 (Gerritsen et al., 1997; Gwack et al., 2001).

### Chromocenters are resistant to VP16, while p65 displays partial activation

The activator VP16, which was found to be weaker than p65 and VPR (Trojanowski et al., 2022), did not induce chromocenter decondensation, H3K27ac and p300 enrichment, or MSR transcription (**Fig. 1C**). This demonstrates that the chromocenter structure maintains a silenced state against a weaker activator. The activation domain of p65 displayed partial activation capabilities. A fraction of cells maintained condensed and silenced chromocenters upon p65 recruitment (“p65 repressed”). In contrast, chromocenters decondensed in other cells and displayed H3K27ac enrichment and p300 recruitment together with MSR transcription (**Fig. 1C**). Upregulated transcription of MSR was never observed in cells with condensed chromocenters, indicating that decondensation is either a requirement for transcription activation or a direct consequence. These results suggest that chromocenters are somewhat protected from activation by transcription factors, with more potent activators overcoming this protective layer.

### HP1α and H3K9me3 remain at chromatin upon decondensation and activation

Next, we wanted to assess how HP1α is distributed at chromocenters. To this end, we performed direct stochastic optical reconstruction microscopy (dSTORM) (Heilemann et al., 2008) of chromocenters. We visualized dCas9 to trace the topology of the associated MSR sequences. Chromocenters displayed a heterogeneous sponge-like chromatin substructure, with areas mostly devoid of MSR and regions with high MSR content (**Fig. 2A**). Upon decondensation by VPR, MSRs dispersed into smaller clusters connected by MSRs in some cases. HP1α showed a granular distribution in both chromatin states with enrichments into smaller subclusters in the range of ∼50 nm with a higher coverage around the dCas9 signal and regions primarily devoid of HP1α and MSRs. These assemblies could be structurally similar to the H3K9me3/HP1 nanodomains of 3-10 nucleosomes found in the non-repetitive part of the genome (Thorn et al., 2022). The dSTORM results indicate that HP1α localization was mainly driven by direct binding to chromatin, but the protein also formed additional, potentially chromatin-independent, subclusters on the scale of ∼100 nm.

**Figure 2.**
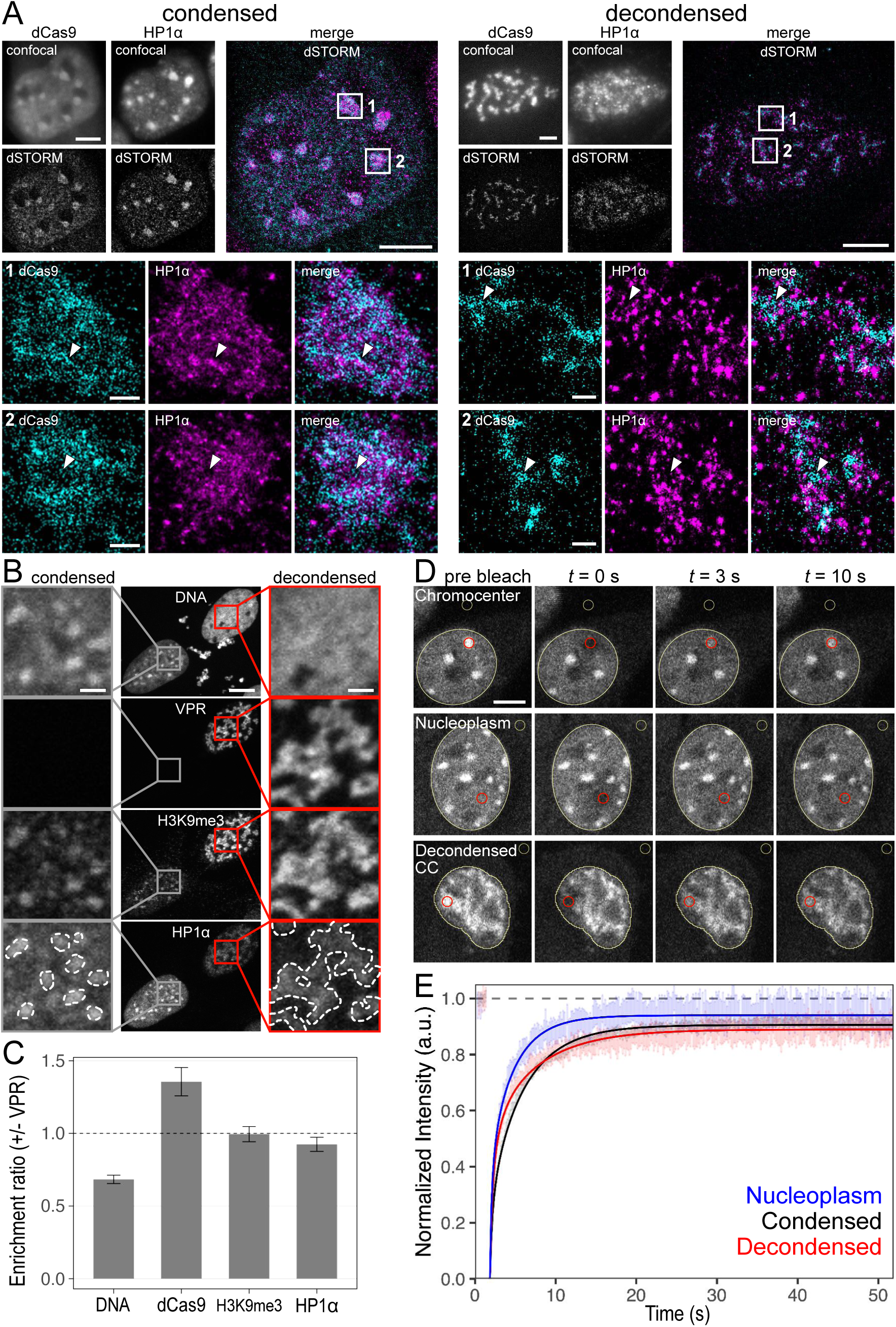
Localization and dynamics of HP1α at chromatin upon decondensation. (**A**) dSTORM images of fixed iMEF cells with dCas9-GFP visualized with an Alexa 488 conjugated GFP-nanobody and endogenous HP1α stained with an Alexa 647 conjugated secondary antibody. Arrows indicate areas of higher HP1α density (condensed 1), regions devoid of dCas9 or HP1α (condensed 2), or smaller clusters connected by MSR (decondensed 1+2). Scale bars, 5 µm (overview) and 0.5 µm (zoom). (**B**) iMEF images of HP1α and H3K9me3 at condensed and decondensed chromocenters. The dotted lines exemplify the chromocenter area used for quantification. Scale bars, 10 µm (overview) and 2 µm (zoom). (**C**) Enrichment ratio between condensed and decondensed chromocenters in iMEFs. The ratio is calculated by dividing the mean enrichment of cells with dCas9-GFP-VPR (decondensed) by the mean enrichment of cells with dCas9-GFP (condensed). The enrichment (per nucleus) was calculated as the mean intensity ratio between chromocenters/nucleoplasm for DNA (DAPI), dCas9 (GFP), H3K9me3 (IF), and HP1α. Error bars represent the standard error of the mean enrichment. Number of cells: dCas9-GFP (condensed) = 21, dCas9-GFP-VPR (decondensed) = 13. (**D**) Exemplary FRAP image series of endogenously GFP-tagged HP1α in 3T3 cells. Cells were transfected with dCas9-tdTomato or dCas9-tdTomato-VPR, respectively, and analysis was started ∼30 h after transfection. The nuclear area is marked by a white outline with red circles delineating bleach spots and white circles indicating the background areas. Scale bar, 5 µm. (**E**) FRAP recovery curves with a fit to the data with a reaction-diffusion model yielding the parameters given in **Supplementary Table S1**.

Next, we investigated what happens to HP1α and H3K9me3 upon decondensation of chromocenters and transcription activation of MSR by VPR. Surprisingly, both HP1α and H3K9me3 retained a strong signal at decondensed chromocenters (**Fig. 2B**). Comparing the mean enrichment of signal inside the chromocenter area between nuclei with condensed and decondensed chromocenters confirmed that H3K9me3 and HP1α enrichment was mostly unaffected (**Fig. 2C**). We conclude that chromocenter decondensation occurs independent of HP1α. Furthermore, HP1α and H3K9me3 can coexist with transcription and H3K27ac, which emerged upon VPR binding (**Fig. 1C**).

### HP1α binding to chromocenters is unaffected by decondensation and transcription

We next examined if the binding dynamics of HP1α at chromocenters were affected by decondensation and transcription induced by VPR. To this end, we performed fluorescence recovery after photobleaching (FRAP) of endogenously tagged HP1α in 3T3 cells transfected with dCas9-tdTomato or dCas9-tdTomato-VPR to measure HP1α mobility at condensed vs. decondensed chromocenters and in the nucleoplasm (**Fig. 2E**). HP1α mobility and chromatin interactions were quantitated by fitting the FRAP curves to a reaction-diffusion model as described previously (Muller et al., 2009; Muller-Ott et al., 2014) (**Supplementary Table S1**). An excellent fit to the data was obtained that yielded (i) an apparent diffusion coefficient *D*app, reflecting the free mobility together with transient binding, (ii) the dissociation rate constant *k*off that describes more long-lived binding interactions, and (iii) the fraction *f*im of protein that was immobile.

Bleached HP1α showed a fast recovery, as observed previously (Muller et al., 2009; Muller-Ott et al., 2014). Recovery curves and values for *k*off and *f*im at condensed and decondensed chromocenters were similar. The only discernable difference was an increase of ∼20% in the diffusive free fraction at the expense of the fraction bound with *k*off at decondensed chromocenters. We conclude that (i) HP1α chromatin binding is mainly unaffected by decondensation and transcription, (ii) HP1α and H3K9me3 are retained on chromatin upon decondensation or activation of transcription, and (iii) HP1α and the repressive mark H3K9me3 can coexist with activating chromatin marks like H3K27ac.

### HP1α can repress transcriptional activation at short time scales

To elucidate the relation between HP1α binding and transcription, transcription activation in the presence of HP1α was further dissected with the U2OS 2-6-3 reporter cell line. Cells were transfected with plasmids encoding rTetR-GFP-activator (VP16, p65, VPR) and GFP-binding protein (GBP)-HaloTag (control) or GBP-HaloTag-HP1α. In addition, the Krüppel associated box (KRAB) domain (Urrutia, 2003) was used as a reference for a canonical repressor (**Fig. 3A**). The GBP-GFP interaction creates a complex of activator and repressor (or the control) that binds to the *tet*O sites of the reporter array upon doxycycline addition. The bulk readout of the reporter transcript was measured by RT-qPCR, while RNA-FISH was applied to analyze transcription in single cells. HP1α repressed transcription activation after 24 h of recruitment to the locus as measured via RT-qPCR of VP16 by 6.9-fold and of p65 by 3.3-fold. At the same time, the effect of HP1α on the potent synthetic activator VPR was marginal (**Fig. 3B**). Repression by HP1α was accompanied by a reduction of RNA polymerase II (RNAP II) and its serine 5 phosphorylated (S5p) form (**Supplementary Fig. S2**). The RNA-FISH analyses in single cells showed that repression occurred by complete silencing of transcription at the locus or by attenuation, where transcription is ongoing at a reduced level (**Fig. 3D**). The fraction of cells silenced upon HP1α recruitment was dependent on the activator type and highest for VP16. At the same time, changes for p65 were moderate and hardly detectable for VPR (**Fig. 3C**). Quantification by RNA- FISH corroborated the RT-qPCR results. It displayed a clear HP1α-mediated reduction for VP16 and p65, while only a slight difference was observed with VPR (**Fig. 3E**).

**Figure 3.**
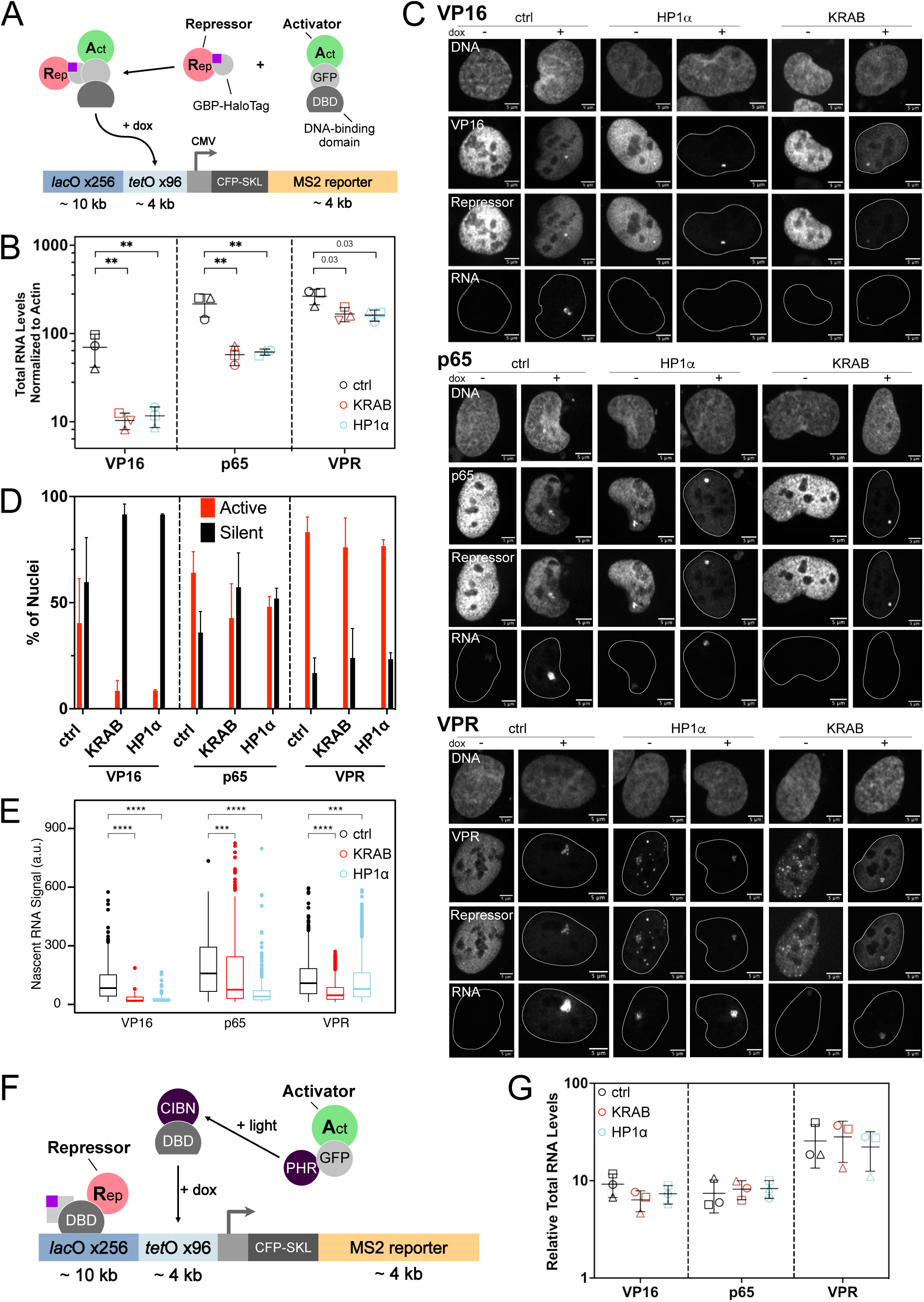
Repression of reporter array transcription by HP1α at promoter-proximal sites. Data refer to measurements at 24 h after transfection. (**A**) Activators (Act) are co-recruited with repressors (Rep) HP1α or KRAB to the reporter locus in the U2OS 2-6-3 cell line in the presence of doxycycline for 24 h. (**B**) Quantification of total RNA levels by RT-qPCR at 24 h post-doxycycline addition. Mean and SD of fold-change induction normalized to beta-actin mRNA multiplied by 1000 (n = 3, shapes represent replicates; see **Methods** for details). One-way ANOVA comparing control to HP1α or KRAB per activator, *p < 0.05, **p < 0.01. (**C**) Confocal microscopy images showing co-recruitment of activator and HP1α or KRAB and its effect on transcription, as detected by RNA-FISH 24 h post-doxycycline addition. Scale bar, 5 µm. (**D**) Fraction of nuclei classified as active or silent based on the presence or absence of transcripts as detected by RNA-FISH 24 h post-doxycycline addition, respectively. Bar: mean, error bars: minimum and maximum of two biological replicates. (**E**) Nascent RNA levels were determined for nuclei classified as active, n = 58- 671, per combination of activator and HP1α, KRAB or control across two biological replicates. Wilcoxon test, ***p < 0.001, ****p < 0.0001. (**F**) HP1α or KRAB was recruited to *lac*O repeats (approx. 4 kb away from the promoter) for 48 h, followed by light and doxycycline-mediated targeting of individual activators to *tet*O repeats for 6 h. (**G**) Quantification of total RNA levels by RT-qPCR. Mean and SD of fold-change induction normalized to beta-actin mRNA and compared to the no doxycycline control (n = 3). Ordinary one-way ANOVA comparing control to HP1α or KRAB per activator, p > 0.05.

To assess whether the repressive effect of HP1α relied on promoter proximity, we transfected cells with HaloTag-LacI-HP1α, CIBN-rTetR and PHR-GFP-activator constructs. HP1α was recruited to the *lac*O sites at a 4 kb distance from the promoter for 48 hours before the activator was recruited to the *tet*O sites in the presence of doxycycline and blue light (**Fig. 3F**). In this experimental setting, HP1α exhibited no repressive effect on transcription when bound to the distal *lac*O sites. The dependence on promoter proximity for the repressive effect of HP1α was consistent across bulk and single-cell experiments (**Fig. 3G**, **Supplementary Fig. S4**).

### HP1α protects chromocenters from transcription activation and decondensation

Next, we wanted to elucidate how the repressive effect of HP1α contributes to chromocenter silencing and resistance against activators. To this end, we recruited the three activators to chromocenters and compared wild-type iMEF cells to a *Suv39h* dn cell line. In the latter cell line, chromocenters are depleted of H3K9me3 and HP1α (**Fig. 1A**). Perturbation by VPR in *Suv39h* dn cells resulted in the same phenotype as seen in wild-type cells (**Fig. 4A**, **Fig. 1C**). Chromocenters had fallen apart into substructures that were enriched in H3K27ac, were in vicinity of p300 and showed transcription of MSR. Similarly, chromocenters in *Suv39h* dn cells affected by p65 were decondensed and showed H3K27ac and transcription of MSR. This is in contrast to the same perturbation in wild-type cells, which only resulted in a subset of cells with decondensed and activated chromocenters (**Fig. 4B**). In addition, perturbation of chromocenters with VP16 in *Suv39h* dn cells significantly increased the range of observed levels of decondensation and transcription compared to the same perturbation in wild-type cells (**Fig. 4B**).

**Figure 4.**
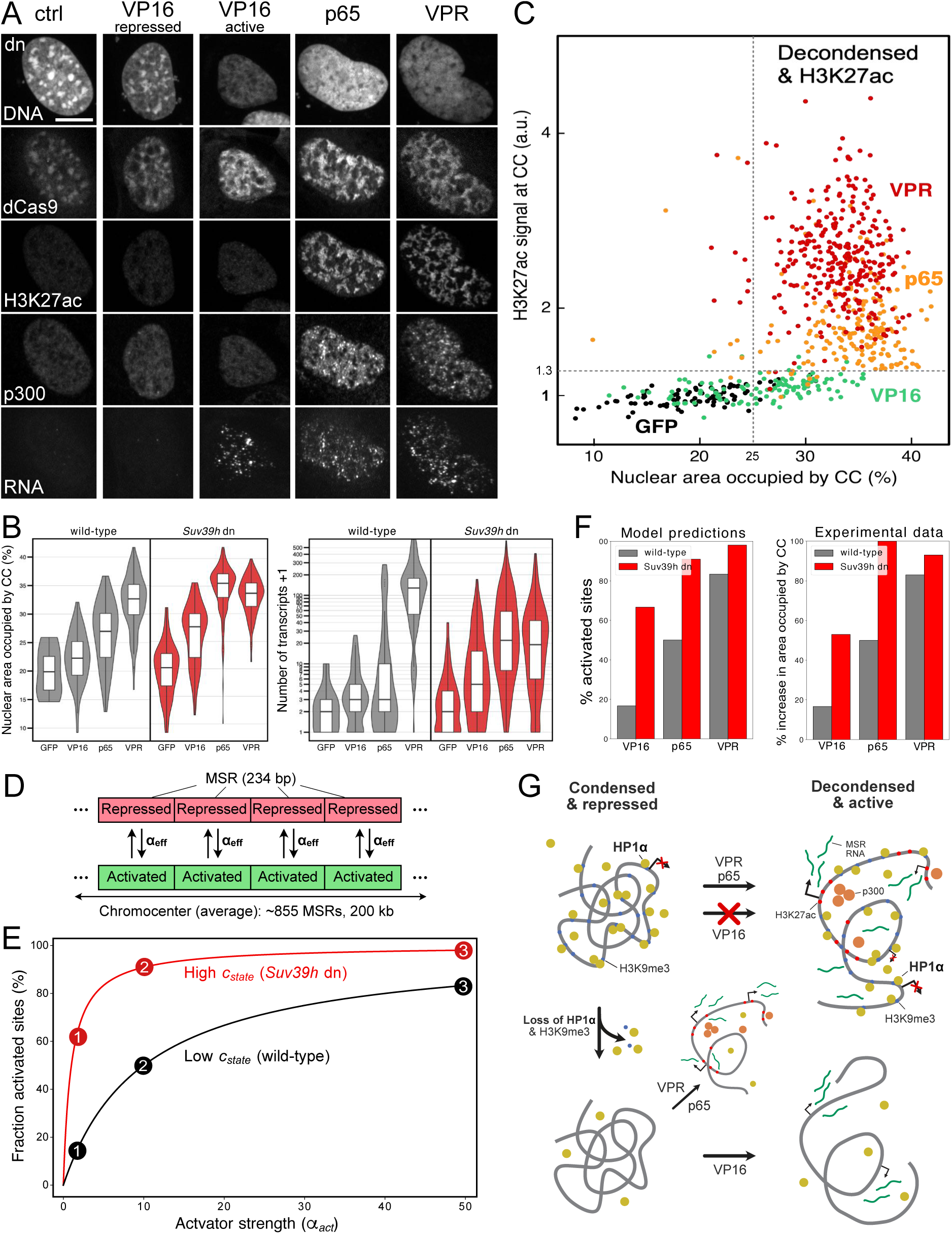
Decondensation of chromocenters and activation of MSR transcription in the presence or absence of HP1α. (**A**) Exemplary confocal images of different activators and their effect on chromocenters in iMEF *Suv39h* dn cells. Scale bar, 10 µm. (**B**) Quantification of decondensation and transcription across conditions. Number of cells for wt: *n* = 52 (VP16), 49 (p65), 109 (VPR), 10 (control). Number of cells for *Suv39h* dn: *n* = 112 (VP16), 188 (p65), (306 VPR), 91 (control). (**C**) Scatterplot of H3K27ac enrichment at chromocenters vs decondensation measured in % of occupied nuclear area. (**D**) Scheme of the 1D lattice model for decondensation of chromocenters. Each repeat was described as an identical and independent unit that can adopt a repressed or activated state. The effective activation rate α*_eff_* is calculated from the chromatin state of each unit and an activator-specific effective activation rate. It describes the transition to the activated state, directly proportional to the degree of chromocenter decondensation. (**E**) Percentage of activated sites over activator strength α*_act_* calculated using the model. Black, wild type; red, *Suv39h* dn with a 10-fold increased *C_state_* value. Numbered circles highlight α*_act_* values and their corresponding % of activated sites representing the activator-induced perturbances in wt and *Suv39h* dn conditions for the same values of α*_act_* (**F**) Left: Model predictions for three fixed values of activator strength. Right: Normalized experimental data from (C) showing the average % increase of nuclear area occupied by chromocenters when induced by VP16, p65, and VPR in wt (grey) and *Suv39h* dn cells (red). (**G**) Model summarizing heterochromatin activation and the involvement of HP1α in it. The repressive ability of HP1α prevents VP16 from activating MSR transcription, and no decondensation is observed. VPR and p65 can overcome this protective layer to induce transcription. (Bottom) Upon loss of HP1α and H3K9me3, VP16 can activate transcription and induce decondensation without significant changes in H3K27ac levels.

While some VP16 cells retained condensed and silenced chromocenters, others now showed decondensed and active chromocenters (**Fig. 4A**). Overall, chromocenters in *Suv39h* dn cells were more susceptible to decondensation and transcription activation by p65 and VP16. We suspect that VPR already induced the maximum perturbance possible for chromocenters in wild-type cells. Therefore, the effect is not further amplified when the repressive network of HP1α is missing in *Suv39h* dn cells. Quantifications of decondensation and transcription across all conditions confirmed these trends (**Fig. 4B**). Interestingly, a small fraction of roughly 2% of *Suv39h* dn cells showed decondensed chromocenters when p65 was recruiting but lacked MSR transcripts (**Supplementary Fig. S5**). This suggests that decondensation alone is not sufficient to induce transcription but accompanies the activation process.

### Loss of HP1α and H3K9me3 facilitates activation of VP16 without enriching H3K27ac

As described above, VP16 was sufficient to decondense chromocenters in *Suv39h* dn cells to some extent. Interestingly, this was achieved without H3K27ac or p300, as we did not observe enrichment of either in any of the VP16 conditions tested (**Fig. 4A**). This is further illustrated by comparing H3K27ac enrichment and level of decondensation across wild-type and *Suv39h* dn (**Fig. 4C**). Here, p65 and VPR cells had up to 3 or 4 times higher H3K27ac enrichment compared to the control respectively. However, VP16-perturbed chromocenters showed H3K27ac levels similar to the control sample, even at high decondensation levels. This indicates that acetylation is not required for decondensation of chromocenters to occur. Therefore, we suggest that other mechanisms can contribute to chromocenter decondensation. This might differ depending on the activator used but likely involves the recruitment of chromatin remodelers or the mediator complex. Additionally, we observed transcription of MSR in *Suv39h* dn cells treated with VP16 (**Fig. 4A, B**). This shows that H3K27ac is neither required for transcription activation nor deposited due to ongoing transcription in this system.

### A 1D lattice ligand binding model rationalizes chromocenter activation data

HP1α has a repressive activity, counteracting the three different activators to a varying extent (**Fig. 3**). To explain our findings quantitatively, we modeled a chromocenter as a 1D lattice of 855 identical and independent units, with each representing one 234 bp long MSR repeat for an average chromocenter size of ∼200 kb (**Fig. 4D, Supplementary Table S2**). In this model, chromatin adopts a compacted state (independent of H3K9me3/HP1α) that decondenses due to histone acetylation and/or RNA synthesis upon induction of transcription (Gibson et al., 2019; Gorisch et al., 2005; Toth et al., 2004). Thus, the decondensation of chromocenter substructures emerges from the transition of repeat units from the silenced state, where they tend to stick together to a non-interacting/repelling activated state. The degree of decondensation of the chromatin fiber was taken to be directly proportional to the number of activated units. The activation reaction via dCas9-mediated targeting of VP16, p65 and VPR to MSRs is described as one activation rate α_*eff*_ that depends on the specific activator rate as well as the pre-existing chromatin state – wild-type HP1α/H3K9me3 levels or *Suv39h* dn cells depleted of HP1α/H3K9me3. The silencing activity of HP1α was estimated to be a 10-fold reduction of the activation rate based on the repressive activity measured for co-recruitment of HP1α with VP16 as a lower bound (**Fig. 3B**), which is represented by the value of the *C_state_* parameter in the model. **Fig. 4E** depicts the number of activated binding sites as a function of the activator strength α_*act*_for either wild-type or *Suv39h* dn cells with *c_state_(wt) = 10 × c_state_*(dn) chromatin. We observed a gradual increase in the nuclear area occupied by chromocenters induced by VP16, p65, and VPR in wild-type cells (**Fig. 4F**). Accordingly, we chose *α_act_* values that recapitulated similar gradual increases in the fraction of activated binding sites for VP16, p65, and VPR (**Fig. 4E, F**) predicted by our model.

Our model predicts a similar pattern in *Suv39h* dn cells, consistent with experimental observations (**Fig. 4F**). VP16 and p65 induce a substantial increase in the fraction of activated repeats in *Suv39h* dn cells. In contrast, VPR induces only a minor increase (**Fig. 4F**). This trend is mirrored in the experiments regarding the increase in the nuclear area occupied by chromocenters under both conditions. Thus, the differential response of wild-type and *Suv39h* dn chromocenters to the recruitment of the three activators concerning decondensation is quantitatively described by this 1D lattice model of independent MSR units. HP1α and H3K9me3 would inhibit the transition from the silenced to the transcribed MSR state with a ∼10-fold reduction of the activation rate. Still, they are not essential for chromatin compaction since their depletion in *Suv39h* dn cells is not sufficient to induce chromocenter decondensation (**Fig. 1A**, **Fig. 4B**). These relations are depicted together with the different activation pathways in **Fig. 4G**.

## DISCUSSION

We dissected the role of HP1α in maintaining the repressive environment of mouse pericentric heterochromatin and protecting against transcriptional activation. Our findings elucidate how chromatin compaction, decondensation, and transcriptional regulation are interlinked. Activation by VP16, p65, and VPR disrupted the chromocenters’ structural integrity and MSR silencing capacity in dependence on the activator strength. HP1α remained stably bound and H3K9me3 persisted at chromocenters on the 24-36 h time scale studied in our experiment, even after decondensation and transcriptional activation by VPR. This challenges the notion that these marks have to be removed for activation. Instead, it suggests that HP1α and H3K9me3 can coexist with active chromatin marks like H3K27ac and during MSR transcription of chromocenters. In *Suv39h* dn cells lacking H3K9me3 and HP1α, chromocenters were more susceptible to decondensation and induction of transcription by VP16 and p65. This underscores the repressive function of HP1α and H3K9me3, which both form a protective barrier against spurious activation.

The repressive activity of HP1α was further dissected using a reporter array that allowed simultaneous recruitment of HP1α (or other repressors) together with the different activators studied here. In this system, we recorded the silencing and attenuation activity of HP1α, i.e., absent vs. reduced transcription in a given cell on the 24-36 h time scale. This process could involve an HP1α-mediated inhibition of RNAP II binding in line with findings for HP1 family proteins *in vitro* and fission yeast (Fischer et al., 2009; Smallwood et al., 2008). HP1α repression was most pronounced for the weaker activator VP16, with p65 and VPR being much less affected and requiring promoter-proximal binding. These findings are consistent with the difference between VP16 activation in wild-type vs. *Suv39h* dn chromocenters. At a 4-10 kb distance to the promoter, HP1α no longer exhibited a repressive effect on transcription, consistent with previous studies showing that H3K9me3 and other histone modifications occurred within a ±4 kb window around a nucleation site (Erdel et al., 2013; Muller-Ott et al., 2014). This result also agrees with a study that showed efficient repression by HP1 at 1-2 kb upstream of a reporter promoter but not at 6.7 kb distance (van der Vlag et al., 2000). Thus, HP1α has a localized repressive activity, which aligns with the co-existence of repressive and activating chromatin marks observed during the activation of chromocenters.

Recruitment of the activators VPR and p65 to MSRs was accompanied by a significant increase of H3K27ac levels, reflecting their capability to directly interact with co-activators like p300 (Trojanowski et al., 2022). High histone acetylation induces an open conformation of the nucleosome chain and drives chromatin decondensation on the µm scale in the cell (Fukai et al., 2024; Gorisch et al., 2005; Otterstrom et al., 2019; Toth et al., 2004). In contrast, VP16-mediated chromocenter transcription in *Suv39h* dn cells was associated with chromatin decondensation but occurred without enrichment of H3K27ac or p300. This observation suggests an alternative decondensation mechanism potentially mediated by the nascent MSR transcripts arising from a previously reported activity of RNA to keep chromatin open (Caudron-Herger et al., 2011), which has been demonstrated for LINE1 repeat RNA (Dueva et al., 2019). Thus, histone hyperacetylation and RNA synthesis could open chromatin structure through a converging mechanism: disrupting nucleosome interactions by neutralizing positive charges on histone tails (Dueva et al., 2019; Tse et al., 1998). The resulting chromocenter decondensation was consistently present when MSR transcripts were detected. However, decondensation alone was insufficient to induce transcription, as inferred from p65 recruitment in *Suv39h* dn cells.

Our data on heterochromatin activation can be quantitatively described with a 1D lattice binding model. It represents chromocenters as a chain of repetitive 234 bp long independent units, each carrying one nucleosome (Packiaraj and Thakur, 2023). Such a model would also be compatible with defining larger units that comprise multiple repeat sequences. These units could represent the ∼50 nm HP1α subclusters observed in the granular distribution on our dSTORM images. Interestingly, similarly-sized H3K9me2/3 and HP1-containing heterochromatin nanodomains or “nucleosome clutches” of 3-10 nucleosomes are abundant in the mammalian epigenome (Thorn et al., 2022). It thus appears conceivable that large H3K9me3 domains on the 0.1-1 MB scale in chromocenters arise from the assembly of repeating small independent units. In our model, the MSR units can transition between repressed and activated states depending on the activation rate of the recruited activator and the presence of HP1α/H3K9me3. Compaction is driven by nucleosome interactions between units in the silenced state, which are lost upon histone acetylation or RNA accumulation. This is described by a direct correlation between units in the active state and decondensation, which aligns with the measured changes in the nuclear area occupied by chromocenters.

When parameterized with the experimental data, our model predicts that weak transcriptional activators like VP16 can only decondense chromocenters depleted of H3K9me3 and HP1α, as in the *Suv39h* dn cells. This behavior emerges from a classical ligand binding framework without invoking a previously proposed liquid-liquid phase separation (LLPS) of HP1α (Keenen et al., 2021; Larson et al., 2017; Strom et al., 2017; Tortora et al., 2023). We find no evidence that the disintegration of chromocenters upon activation affects higher-order HP1α assembly. Instead, the transition into a transcriptionally active state occurred with no apparent effect on the local HP1α environment: (i) Super-resolution dSTORM microscopy revealed that HP1α displays a granular distribution at both condensed and decondensed chromocenters, consistent with persistent binding to specific sites. (ii) HP1α mobility and chromatin interactions are mainly unaffected by chromocenter decondensation and transcriptional activation. (iii) The short-term repressive effects of HP1α in the reporter array system remained localized and dependent on promoter proximity.

A silencing mechanism for pericentric heterochromatin, where a localized repressive activity of HP1α/H3K9me3 is present at independent repeats (or repeat clusters) together with transcriptionally active sites, could also apply to the thousands of H3K9me2/3 and HP1-containing heterochromatin nanodomains that are present in the mammalian epigenome (Thorn et al., 2022). A key question will be understanding how cells precisely control the transition of these local barriers against transcriptional activation between silenced and activated chromatin states. Such mechanisms must be sufficiently stable to maintain epigenetic memory while remaining responsive enough to allow controlled gene activation during development and cellular differentiation. The quantitative framework presented here provides a foundation for dissecting these fundamental aspects of chromatin regulation.

## DATA AND CODE AVAILABILITY

The software for image and FRAP analysis is available via GitHub at https://github.com/RippeLab/. Additional information required to reanalyze the data reported in this paper is available from the lead contact upon request.

## ACKNOWLEDGMENTS

We thank Anne Rademacher for discussion and the BioQuant Advanced Biological Screening Facility and DKFZ Light Microscopy Core Facility for their help. This project was supported by grant RI1283/16-2 in the Priority Program 2191 “Molecular Mechanisms of Functional Phase Separation” of the Deutsche Forschungsgemeinschaft (DFG) and by the Cooperation Program in Cancer Research of the German Cancer Research Center (DKFZ) and Israel’s Ministry of Science, Technology and Space (MOST) via grant Ca215. JFN was supported by the inter-institutional postdoc program of the Innovation Campus Health + Life Science Alliance Heidelberg Mannheim (MWK32-7531-32/7/2). Data storage at SDS@hd was funded by the Ministry of Science, Research and the Arts Baden-Württemberg (MWK) and the DFG through grants INST 35/1314-1 FUGG and INST 35/1503-1 FUGG.

## AUTHOR CONTRIBUTIONS

Study design: KR, RW. Data acquisition: RW, AU, JFN. Analysis of data: RW, AU, JFN, KR; Technical assistance: LF, CK; Drafting of manuscript: KR, RW. Manuscript reviewing: all authors. Supervision and coordination: KR

## METHODS

**KEY RESOURCES TABLE**

| REAGENT or RESOURCE | SOURCE | IDENTIFIER |
| --- | --- | --- |
| Antibodies |  |  |
| Mouse monoclonal anti-HP1α | Euromedex | 2HP-1H5-AS |
| Rabbit polyclonal anti-H3K9me3 | abcam | ab8898 |
| Rabbit polyclonal anti-H3K27ac | abcam | ab4729 |
| Mouse monoclonal anti-KAT3B/p300 | abcam | ab54984 |
| GFP-nanobody Alexa 488 | Chromotek | gb2AF488 |
| Rabbit polyclonal anti-RNAPII CTD | abcam | ab26721 |
| Rat monoclonal anti-RNAPII Ser5p | ActiveMotif | 61085 |
| Goat polyclonal anti-rat Alexa 568 | Thermo Fisher Scientific | A-11077 |
| Goat polyclonal anti-rabbit Alexa 568 | Thermo Fisher Scientific | A-11036 |
| Goat polyclonal anti-rabbit Alexa 647 | Thermo Fisher Scientific | A-21244 |
| Goat polyclonal anti-mouse Alexa 750 | Thermo Fisher Scientific | A-21037 |
| Goat polyclonal anti-rabbit Alexa 750 | Thermo Fisher Scientific | A-21039 |
| Biological Samples |  |  |
| iMEF wt cell line | (Muller et al., 2009) | iMEF-wt-cx669 |
| *Suv39h* double-null iMEF cell line | (Peters et al., 2001) | iMEF-*Suv39* dn |
| 3T3 GFP-HP1α cell line | (Muller et al., 2009) | 3T3-HP1α-cx361 |
| U2OS cell line with reporter array | (Janicki et al., 2004) | U2OS 2-6-3 |
| Plasmids |  |  |
| psgRNA-2xMS2-mSat | (Erdel et al., 2020) | Addgene #181908 |
| pdCas9-GFP | (Erdel et al., 2020) | Addgene #181906 |
| pdCas9-GFP-VP16 | (Trojanowski et al., 2022) | Addgene #183923 |
| pdCas9-GFP-p65 | This study | cx1618 |
| pdCas9-GFP-VPR | (Erdel et al., 2020) | Addgene #181907 |
| pdCas9-tdTomato | This study | cx1184 |
| pdCas9-tdTomato-VPR | This study | cx1580 |
| rTetR-GFP | (Trojanowski et al., 2022) | Addgene #183916 |
| rTetR-GFP-VP16 | (Trojanowski et al., 2022) | Addgene #183917 |
| rTetR-GFP-p65 | This study | cx1571 |
| rTetR-GFP-VPR | (Trojanowski et al., 2022) | Addgene #183918 |
| HaloTag-GBP | (Trojanowski et al., 2022) | cx1574 |
| GBP-HaloTag-mHP1α | This study | cx1671 |
| HaloTag-GBP-KRAB | This study | cx1664 |
| HaloTag-LacI | This study | cx1691 |
| HaloTag-LacI-mHP1α | This study | cx1689 |
| HaloTag-LacI-KRAB | This study | cx1688 |
| CIBN-rTetR | (Trojanowski et al., 2022) | Addgene #183919 |
| PHR-GFP | (Trojanowski et al., 2022) | Addgene #183926 |
| PHR-GFP-VP16 | (Trojanowski et al., 2022) | Addgene #183927 |
| PHR-GFP-p65 | (Trojanowski et al., 2022) | Addgene #183929 |
| PHR-GFP-VPR | (Trojanowski et al., 2022) | Addgene #183928 |
| Critical Commercial Assays |  |  |
| X-tremeGENE 9 DNA transfection reagent | Roche | 6365787001 |
| SYBR Green PCR master mix | Applied Biosystems | 4308155 |
| RNAscope Fluorescent Multiplex Kit (v1) | ACD Bio | 320293 |
| RNAscope Probe-MmsatelliteDNA-sense | ACD Bio | 475521 |
| RNAscope Probe-MmsatelliteDNA-sense-C2 | ACD Bio | 475521-C2 |
| Deposited data |  |  |
| Microscopy source images | BioImage Archive<br>https://www.ebi.ac.uk/bio-image-archive/ | Submission in progress |
| Software and Algorithms |  |  |
| Fusion software 2.4 | Andor Technology | https://andor.oxinst.com/downloads/view/fusion-release-2.4 |
| FIJI/ImageJ | https://imagej.net/ | https://www.doi.org/10.1038/nmeth.2089 |
| ThunderSTORM | https://github.com/zitmen/thunderstorm | https://doi.org/10.1093/bioinformatics/btu202 |
| TurboReg | https://bigwww.epfl.ch/thevenaz/turboreg/ | https://bigwww.epfl.ch/publications/thevenaz9801.html |
| Cellpose 2.2.3 | (Stringer et al., 2021) | https://pypi.org/project/cellpose/ |
| Ilastik 1.4.0 | https://www.ilastik.org/download | https://doi.org/10.1038/s41592-019-0582-9 |
| R software environment 4.2.2 | R Core Team, 2022 | https://www.r-project.org/ |
| EBImage R package | https://bioconductor.org/packages/release/bioc/html/EBImage.html | https://doi.org/10.1093/bioinformatics/btq046 |

### Lead contact

Further information and requests for resources and reagents should be directed to the corresponding author, Karsten Rippe.

### Materials availability

Plasmids generated in this study will be available from Addgene at https://www.addgene.org/Karsten_Rippe/ upon publication.

### Data and code availability

The image analysis software will be available for custom scripts via the GitHub repository https://github.com/RippeLab/ and for other software via the sources listed in the Key Resources Table. Additional information required to reanalyze the data reported in this paper is available from the lead contact upon request.

## METHOD DETAILS

### Cell Lines

The iMEF Suv39H1/H2 double null and NIH 3T3 GFP-HP1α cell lines have been described previously (Muller et al., 2009; Peters et al., 2001). The U2OS 2-6-3 female osteosarcoma cell line with a *lac*O/*tet*O reporter gene array (Janicki et al., 2004) and the different approaches for recruiting proteins to this reporter (Trojanowski et al., 2022) have been described previously. All cell lines were tested for the absence of mycoplasma with the VenorGeM Advance kit (Minerva Biolabs) and human cell lines were authenticated using single nucleotide polymorphism profiling (Multiplexion).

### Plasmids

Protein constructs were expressed with a CMV promoter using pEGFP-C1/N1 (Clontech) (enhanced GFP, referred to here as GFP) or pcDNA3.1 (Invitrogen) vector backbones. A U6 promotor-driven expression vector containing the sgRNA with MSR-targeting sequence (GGG CAA GAA AAC TGA AAA TCA) was described previously (Erdel et al., 2020).

### Cell culture

Cells were cultured at 37°C and 5% CO2 in DMEM supplemented with 10% fetal calf serum (PAN), 2 mM L-glutamine (PAN), 1% penicillin/streptomycin (PAN), and 1 g/l glucose for U2OS or 4.5 g/l glucose for iMEF and 3T3 cells. For microscopy analysis, U2OS cells were seeded onto 8-well chambered coverglass slides (Nunc Lab-Tek, Thermo Fisher Scientific) at a density of 3 x 10^4^ cells per well and 7.5 x 10^3^ cells per well for experiments with 24 h and 48 h until fixation respectively. For RT-qPCR, 2-3 x 10^5^ cells or 7.5 x 10^4^ cells were seeded in 6-well plates, respectively. iMEF and 3T3 cells were seeded onto 12 mm coverslips (borosilicate #1, Thermo Fisher Scientific) at a density of 3 x 10^4^ cells per well. One day after seeding, cells were transfected with X-tremegene 9 DNA transfection reagent (Roche) according to the manufacturer’s protocol. For U2OS cells, 300 ng or 500 ng of total plasmid DNA was used per well for experiments with 48 h and 24 h until fixation, split equally across constructs. For RT-qPCR, transfection reactions were scaled up to 1.8 µg and 2 µg, respectively. For iMEF and 3T3 cells, 1 µg of total plasmid DNA was used per well, split equally across constructs. Cells used for optogenetic experiments were protected from light until the induction time point and then exposed to diffused white light for 6 h until fixation or harvest. Cells transfected with HaloTag plasmids were labeled with 0.2 µM Janelia Fluor 646 Halo-tag ligand (Promega) in the medium for 20 min in the dark. Medium containing 5 µg/ml doxycycline (Sigma-Aldrich) was used to recruit rTetR constructs.

### RNA isolation and RT-qPCR

Cells were harvested after trypsinization using the Nucleospin RNA Plus Kit (Macherey-Nagel, 740955). The isolated RNA was treated for 30 min at 37°C with RQ1 DNase (Promega) according to the manufacturer’s protocol and then purified using the Nucleospin RNA Clean-Up XS kit (Macherey-Nagel). RNA concentration and purity were determined by absorbance measurement. Per sample, 750-1000 ng of RNA was used based on the total yield for cDNA synthesis using the Superscript IV reverse transcriptase protocol (Thermo Fisher Scientific). RT-qPCR was performed in technical triplicates with 2 µl of 1:40-diluted (if 1000 ng total) or 1:30-diluted (if 750 ng total) cDNA per 10 µl reaction using SYBR Green PCR master mix (Applied Biosystems) with a final primer concentration of 500 nM. The following PCR primers (Eurofins Genomics) were used. Human beta-actin forward: 5’-TCC CTG GAG AAG AGC TAC GA-3’, reverse: 5’-AGC ACT GTG TTG GCG TAC AG-3’; CFP-SKL forward: 5’-GTC CGG ACT CAG ATC TCG A-3’, reverse: 5’-TTC AAA GCT TGG ACT GCA GG-3’. RT-qPCR was analyzed by normalizing reporter RNA levels to beta-actin mRNA levels (ΔCT) alone or as a fold change of the control.

### RNA-FISH

Cells were fixed by adding 4% paraformaldehyde (in PBS, Sigma-Aldrich, 252549) to the medium (final concentration 2%) for 5 min, followed by aspiration and additional fixation in 4% paraformaldehyde in PBS for 15 min. For iMEF cells, the RNAscope system (ACD Bio) with RNAscope Fluorescent Multiplex Kit (v1) and catalog probes RNAscope Probe-Mmsatellite-DNA- sense or Probe-Mmsatellite-DNA-sense-C2 were used with diethylpyrocarbonate treated PBS. After washing fixed iMEF cells three times with PBS, they were incubated with 3% hydrogen peroxide for 5 min and washed twice with PBS. Target and amplification probes were then hybridized according to the manufacturer’s protocol. After amplification, target probes were labeled with Atto Fluor 550 using the C1 or C2 detection kit. Cells were stored in PBS overnight before performing immunofluorescence staining as described below, omitting the fixation step. Fixed U2OS cells were washed three times with PBS, permeabilized with 0.5% Triton X-100 for 5 min, and washed 3 times with PBS. An additional RNase treatment was performed with 0.5 mg/ml RNase A (Thermo Fisher Scientific) in PBS for 1 h at 37 °C, followed by PBS washes.

Samples were dehydrated by sequential washes of 70%, 85%, and 100% ethanol for 3 min each. For hybridization, a mix of 100 nM MS2 probe, 2 µg/µl salmon sperm DNA and 80% formamide was incubated for 10 min at 70°C with equal volumes of 10 mM ribonucleoside-vanadyl complex in 2x hybridization buffer (30% w/v dextran sulfate, 2 mg/ml RNase-free bovine serum albumin in 4x RNase-free saline-sodium citrate buffer (SSC) (Invitrogen). Samples were incubated for 2 h at 37 °C with the hybridization mix and washed with 50% formamide in 2x RNase-free SSC for 15 min each, followed by a 2x RNase-free SSC and DEPC-treated PBS wash for 5 min each. Samples were stored and imaged in DEPC-treated PBS. The sequence of the Atto550 labelled MS2 probe (Eurofins Genomics) was 5’-GTC GAC CTG CAG ACA TGG GTG ATC CTC ATG TTT TCT AGG CAA TTA-3’.

### Immunofluorescence

Cells were fixed by adding 4% paraformaldehyde in PBS (Sigma-Aldrich) to the medium at a final concentration of 2% for 5 min, followed by aspiration and additional fixation in 4% paraformaldehyde in PBS for 15 min. Cells were permeabilized with 0.2% Triton X-100 (Merck) in PBS for 12 min and blocked with 10% goat serum (Cell Signaling Technology) in PBS for 1 h. Samples were incubated with 1:500 diluted primary antibodies in 10% goat serum in PBS for 1 h at room temperature, followed by 3 washes with 0.002% NP40 (Sigma-Aldrich) in PBS for 5 min each. Secondary antibodies conjugated with Alexa488/568/647/750 were diluted 1:250 in 10% goat serum in PBS and incubated with the samples for 1 h at room temperature. After 3 washes with PBS, DNA was stained with a 5 µM DAPI solution in PBS for 15 min at room temperature. Coverslips were rinsed in water, dehydrated in 100% ethanol, and mounted with ProLong Diamond Antifade Mountant (Thermo Fisher Scientific). Lab-Tek slides were washed twice for 5 min with PBS and stored in PBS at 4 °C until imaged. Details on antibodies used are given in the **Key Resources Table**.

### Microscopy instrumentation

Fluorescence microscopy imaging and dSTORM (Heilemann et al., 2008) were conducted with an Andor Dragonfly 505 spinning disc microscope equipped with a Nikon Ti2-E inverted microscope and 100x silicone immersion objective (CFI SR HP Apochromat Lambda S 100x, Nikon). Multicolor images were acquired using laser lines at 405 nm (DAPI), 488 nm (GFP), 561 nm (tdTomato, Atto550, JF568), 637 nm (JF646, Alexa647) and 730 nm (Alexa750) for excitation with a quint-band dichroic unit (405, 488, 561, 640, 750 nm) and corresponding emission filters of 445/46 (DAPI), 521/38 (GFP), 571/78 (tdTomato, Atto550), 685/47 nm (JF646, Alexa647) and 700/75 nm (Alexa750). Images were recorded at 16-bit depth with 1024 x 1024 dimensions (pixel size: 0.1205 µm) using an iXon Ultra 888 EM-CCD camera. A Leica TCS SP5 II confocal microscope (Leica) equipped with a 63x Plan-Apochromat oil immersion objective was used for FRAP experiments.

### Image analysis

Images from chromocenter perturbation experiments in iMEF and 3T3 cells were processed and analyzed using the workflow previously described (Frank et al., 2021) and depicted in **Supplementary Fig. S1.** In short, they were imported in ImageJ and projected in z using maximum intensity projection (Schindelin et al., 2012). Cell nuclei segmentation was performed in RStudio using the DAPI signal and an intensity-based thresholding function from the EBImage package (Pau et al., 2010). Chromocenters were segmented with a custom function using the dCas9-GFP signal. The fraction of nuclear area covered by chromocenters was calculated by dividing the total chromocenter mask area by the total nuclear area per cell. Enrichment of a given chromatin feature was calculated for each cell as the ratio of the mean intensities of the chromocenter and nucleoplasm masks and averaged over all cells of a given condition to yield the mean enrichment. Mean enrichments at condensed and decondensed chromocenters were then divided to obtain the enrichment ratio. The number of MSR transcripts was determined via spot calling on the RNAscope signal using the RS-FISH plugin in ImageJ (Bahry et al., 2022) (Sigma = 0.85, Threshold = 0.002, default RANSAC parameters, no background subtraction) and counting the number of spots located inside the nuclear mask.

Images of U2OS cell experiments were imported in ImageJ and projected in z using maximum intensity projection followed by flat-field and chromatic aberration correction. Nuclei were segmented with Cellpose v2.2.3 and the pre-trained model “cyto.” The reporter array was segmented using Ilastik v1.4.0 (Berg et al., 2019) based on the GFP and Janelia Fluor 646 signal for the co-recruitment experiment or based on only the Janelia Fluor 646 signal for the experiment with distal recruitment of the repressor. A probability value was assigned to each segmented reporter array. Images and segmented masks were imported into RStudio using the EBImage package. Images were filtered by removing segmentation artifacts, and quantification of image features was performed using custom R scripts based on the EBImage package. Nucleus masks at image borders were excluded, and nuclear shape features were computed to exclude apoptotic or mitotic nuclei according to a size threshold. A probability threshold was determined, and the predicted reporter array mask from Ilastik was included for analysis. Selected reporter array masks were used to compute RNAP II enrichments by a log2 transformation of the ratio between the mean intensity at the reporter compared to a ring around its boundary. The signal in the ring represents the nucleoplasmic background signal. MS2 RNA signal was quantified using the nuclear mask since RNA would defuse away from the reporter region.

### FRAP analysis

The confocal FRAP experiments were conducted as described previously (Muller-Ott et al., 2014) on the Leica SP5 microscope using the Leica LAF software and bleaching with the argon laser lines (458 nm, 476 nm, 488 nm, 496 nm). Images of 128×128 pixels with a zoom factor of 12.5 corresponding to 155 nm/pixel were recorded at 400 Hz line frequency, resulting in a frame time of 168 ms. For each cell, 10 pre-bleach and 2 bleach frames were recorded with a 1.25 µm diameter circular bleach region. It was placed on the chromocenter (“bleach spot”) or in the background outside the nucleus (“background area”). Subsequently, 400 post-bleach frames were recorded. Images without the two bleach frames were registered using TurboReg with affine and accurate settings using the first pre-bleach frame as the target and the rest of the time course as the source. Registered images were then used to calculate the bleach profile for each condition. To this end, a radial profile was calculated in ImageJ based on the difference in the image of the first post-bleach frame and the last pre-bleach frame. Radial profiles were normalized to the mean intensity of a background region, averaged, interpolated, and fitted using the following constant function with Gaussian edges using R based on (Mueller et al., 2008):

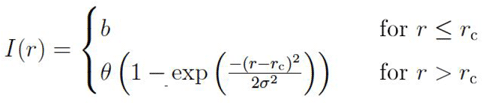

A mean FRAP curve and the standard error of the mean were calculated from all FRAP curves per condition using the first 300 post-bleach frames in R. This mean curve was then fitted in the FREDIS software using a reaction-diffusion model as previously described (Muller et al., 2009; Sprague et al., 2004) but using the radius of the constant (*r_c_*) and the width of the Gaussian (*σ*) as input parameters since the newer version of FREDIS used here also takes the shape of the initial bleach profile into account. FREDIS calculated the diffusion coefficient *D*, the effective diffusion coefficient *D_eff_*, dissociation rate *k_off_*, pseudo on rate *k*_on_*, pseudo affinity *k*_aff_*, free and bound protein fraction, immobile fraction *f_i,_* and the coefficient of determination *R^2^*. Diffusion coefficients of all conditions were averaged and resulted in *D* = 3.26 µm^2^s^-1^. Mean FRAP curves were fitted again in FREDIS using this average diffusion coefficient as a fixed parameter for the reaction-diffusion model, and the final output values were collected in **Fig. 2F**. The resulting fits and the input mean curves were normalized to the bleach depth and mean of the pre-bleach frames and plotted in R.

### dSTORM high-resolution imaging

Cells were transfected with dCas9-GFP-VPR and fixed by adding 4% paraformaldehyde (in PBS, Sigma-Aldrich, 252549) to the medium (final concentration 2%) for 5 min, followed by aspiration and additional fixation in 4% paraformaldehyde in PBS for 15 min. Samples were immunostained with an Alexa 488 conjugated GFP-nanobody (1:500, Chromotek) according to the manufacturer’s protocol, as well as a mouse monoclonal anti-HP1α primary antibody (Euromedex) and a goat polyclonal anti-mouse Alexa 647 conjugated secondary antibody (Thermo Fisher Scientific). Cells were imaged in a reducing oxygen scavenging buffer containing 10% glucose (w/v), 10 mM Tris (pH 8.0), 2x SSC, 37 µg glucose oxidase (Sigma, G2133), 1% (v/v) catalase (Sigma C3515) and 143 mM β-mercaptoethanol. For each experiment, 5000 frames were acquired in widefield mode with a power density illumination lens PD4, full intensities of the 488 nm or 637 nm lasers, and 50 ms exposure time. Image series were analyzed using the Thunderstorm plugin in ImageJ (Ovesny et al., 2014). A B-spline wavelet filter with order = 3 and scale = 2.0 was used for image filtering. The local maximum method approximated the localization of molecules with a peak intensity threshold and 8-neighborhood connectivity. Sub-pixel molecule localization was performed using an integrated Gaussian method with a fitting radius of 4 pixels, a maximum likelihood fitting model, and an initial sigma of 1.5 pixels. dSTORM images were created by averaged shifted histogram rendering with 5-fold magnification and lateral shifts of 2.

### 1D lattice model for chromocenter activation

The model represents a chromocenter as a 1D lattice, with each unit representing one 234 bp long pericentromeric satellite DNA repeat. Each of the units is independent and can be in an activated or repressed state. An average number of 855 MSRs per chromocenter is calculated from a DNA content of 6.4 MB of the pericentric repeats in mouse cells (corresponding to 27,350 MSRs) and an average of 32 DAPI-stained chromocenters per nucleus (**Supplementary Table S2**). In the model, repressed binding sites are assumed to associate with each other, while activated binding sites do not interact with or repel each other. Thus, decondensation of chromocenter substructures emerges from the transition of repeat units from the repressed to the activated state. In experiments, activation is induced by the dCas9-mediated targeting of VP16, p65 and VPR to the satellite repeats for which saturating binding-site occupancy is assumed. Activator binding causes the transition to an activated state via downstream interactions with other co-activators that lead to decondensation (e.g., through acetylation of histones or transcription itself). All these reactions are cast into an effective activation rate α_*eff*_, which depends on the specific activator rate *c_state_* and the pre-existing chromatin state *c_state_*. The latter parameter can be wild-type with corresponding HP1α/H3K9me3 levels or *Suv39h* dn mutated and depleted of HP1α/H3K9me3. We also assume the chromatin fiber decondensation is directly proportional to the number of activated binding sites. For each binding site *C_i_*, the activation equilibrium can be written as:

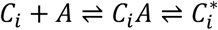

*C_i_* is the *ith* binding site on the chromatin fiber, *C_i_A* is an activator, *C_i_ A* is an activator-bound binding site, and 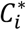 is an activated binding site. At saturating activator concentrations

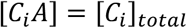

where 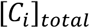 is the total concentration of *ith* binding sites.

In the second step, activator-bound sites *C_i_ A* transition to the activated state 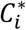. The effective activation rate *α_eff_* is then given by:

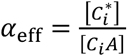

The value of α*_eff_* depends on the chromatin (H3K9me3 and HP1α present or depleted) and the activator’s propensity to activate binding sites. Thus, 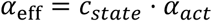 and we yield:

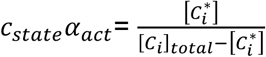

The parameter 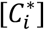 is expressed by:

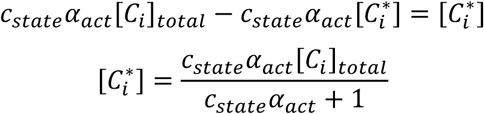

The fraction of activated binding sites θ is given as:

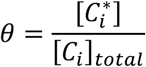

Substituting the expression derived for 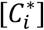, we get:

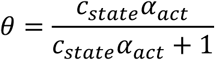

For *N* = 855 independent and equal binding sites, the total number of activated binding sites *n_act_* is:

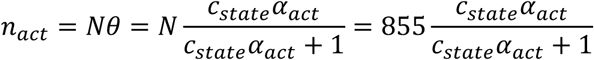

Without loss of generality, we set *c_state_* to be unity for active chromatin (which mimics *Suv39h* dn cells). From our experimental data, we estimated *c_state_* to be reduced 10-fold yielding:

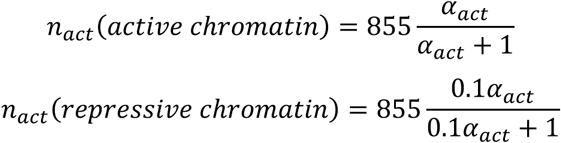

The experimentally observed maximum of the averaged nuclear area occupied by chromocenters (CC) was *comp_max_*= 35% for p65 in *Suv39h* dn, and the minimum was *comp_min_* = 20% in the GFP control in wild-type cells. Therefore, the general formula to calculate the degree of decondensation induced by an activator is:

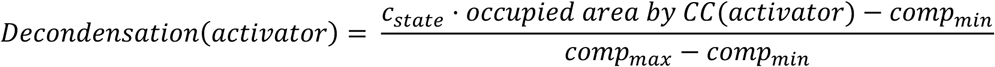

yielding the following decondensation values in wild-type cells:

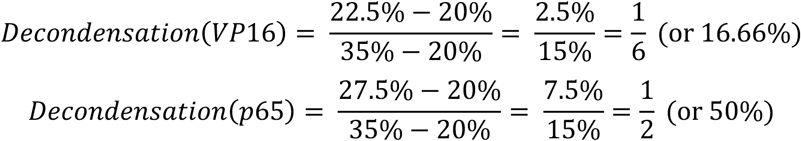

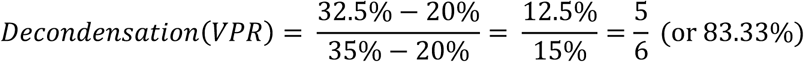

In our model, the average fraction of activated binding sites is directly proportional to the activator-induced decondensation. Accordingly, we can calculate the respective activator strengths (α_act_) by rearranging the above-derived expression for the fraction of activated binding sites θ =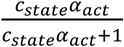 to get:

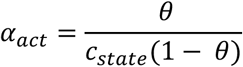

With the activator-specific values for the degree of decondensation as the average fraction of activated binding sites and the *c_state_* value for wild-type cells, we obtain:

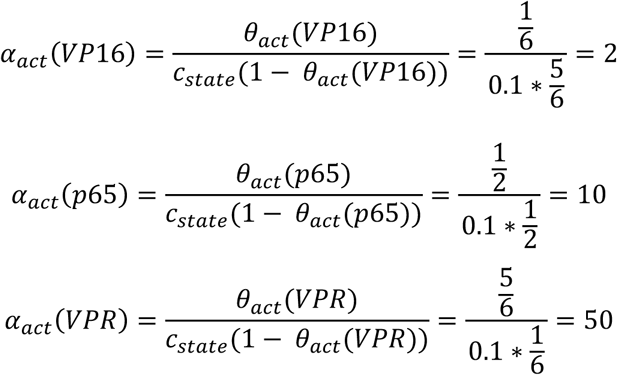

## QUANTIFICATION AND STATISTICAL ANALYSIS

Mean values and 95% confidence intervals (CI) were calculated using a Student’s t-distribution. Pairwise comparisons of the mean for RT-qPCR or relative intensities were conducted using ordinary one-way ANOVA or Wilcoxon’s tests, respectively. To assess the effects of two grouping variables (presence of HP1α for its impact on transcription), a two-way ANOVA (type II) was performed. Error bars represent one standard deviation (s. d.) for RT-qPCR experiments and the standard error of the mean (s. e. m.) as indicated. Axis breaks were introduced in relative intensity and RT-qPCR plots for conditions with values on very different scales or with outliers and are marked by an interruption of the axis. Box plots display the first and third quartile (box), median (bar), data points within the 1.5-fold interquartile range (whiskers), and outliers (points). The number of cells (n) is reported in the figure legends.

