## Supplementary Information for "HP1 binding creates a local barrier against transcription activation and persists during chromatin decondensation"

### SUPPLEMENTARY TABLES

**Table S1. FRAP fit parameters**

| | CC condensed | CC decondensed | Nucleoplasm |
|---|---|---|---|
| $D_{eff}$ (µm² $s^{-1}$) | 1.65 | 2.29 | 2.10 |
| $k_{off}$ ($s^{-1}$) | 0.30 [0.29…0.31] | 0.18 [0.17…0.19] | 0.38 [0.35…0.42] |
| $k^*_{on}$ ($s^{-1}$) | 0.29 [0.28…0.31] | 0.08 [0.07…0.08] | 0.21 [0.19…0.23] |
| $K^*_{eq}$ | 0.98 [0.90…1.02] | 0.43 [0.39…0.46] | 0.55 [0.49…0.61] |
| $f_{free}$ (%) | 50 [49…52] | 70 [68…71] | 64 [62…66] |
| $f_{bound}$ (%) | 50 [48…51] | 30 [29…32] | 36 [34…38] |
| $f_{im}$ (%) | 9 | 11 | 6 |
| n | 32 | 15 | 14 |

**Table S2. Model parameters**

| | |
|---|---|
| Number of repeats [a] | 855 |
| $c_{state}$(repressive chromatin) | 1 |
| $c_{state}$(active chromatin) | 0.1 |
| $\alpha_{act}(VP16)$ | 2 |
| $\alpha_{act}(p65)$ | 10 |
| $\alpha_{act}(VPR)$ | 50 |

[a] The number of repeat units per chromocenter was estimated based on 6.4 Mb of pericentromeric repeats, a major satellite repeat length of 234 bp, and an average of 32 DAPI-stained chromocenters per nucleus (Eck et al., 2012; Muller-Ott et al., 2014).

#### SUPPLEMENTARY FIGURES

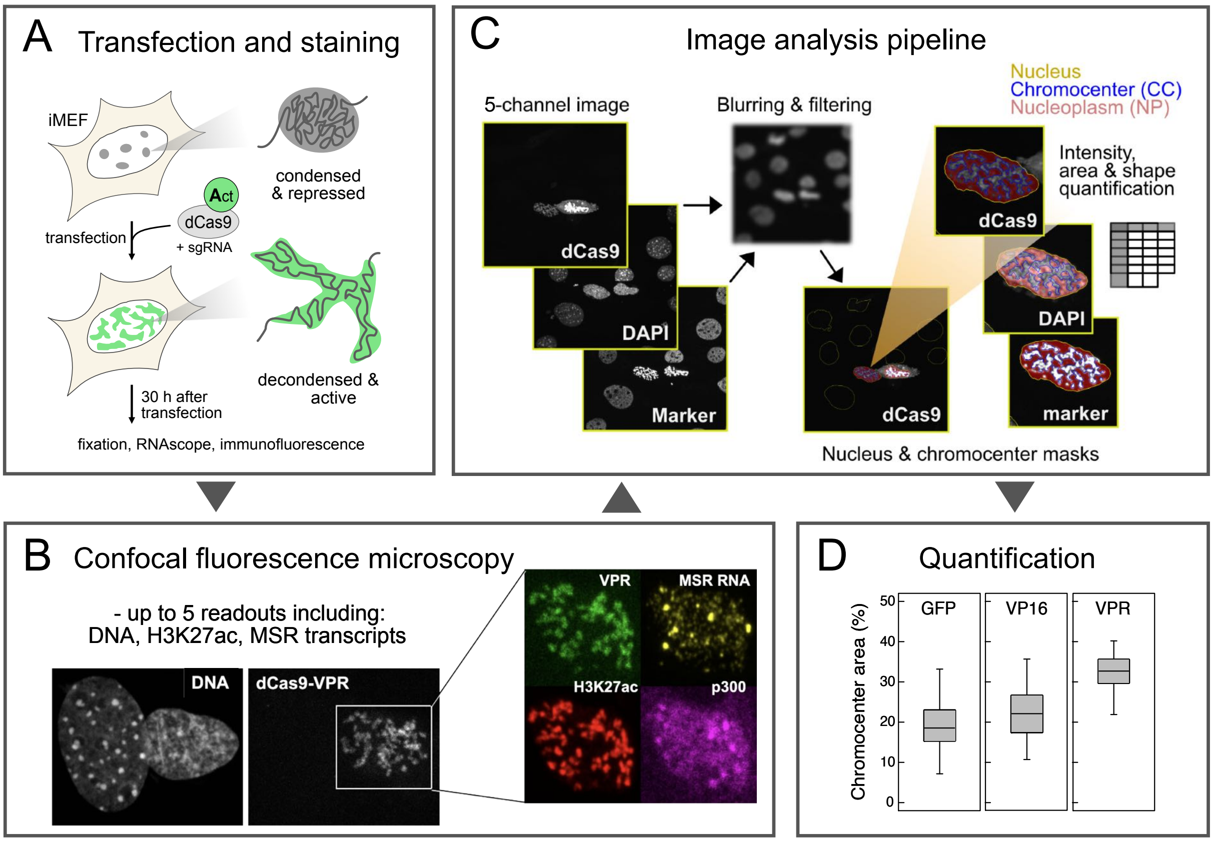


**Figure S1. Image analysis pipeline** for chromocenter decondensation and transcription activation. (**A**) Transfection and staining. iMEFs were transfected with dCas9 and gRNAs targeting MSRs. At 30 h post-transfection, cells were fixed and subjected to RNAscope and immunofluorescence staining. (**B**) Fluorescence microscopy. Images were captured with up to five readouts, including DAPI, dCas9, and specific markers. (**C**) Adapted from Frank *et al.*, 2021. Image analysis pipeline. Images underwent blurring to enhance segmentation accuracy and filtering to exclude artifacts. Using these processed images, masks were generated for the nucleus and chromocenters by using thresholding segmentation. Intensity and area quantifications were performed, distinguishing the nucleus (yellow), chromocenter (blue), and nucleoplasm (purple). (**D**) Quantification. The chromocenter area percentage was quantified across different conditions (GFP, VP16, VPR). Box plots display the chromocenter area percentages, indicating the level of decondensation of chromocenters in each condition.

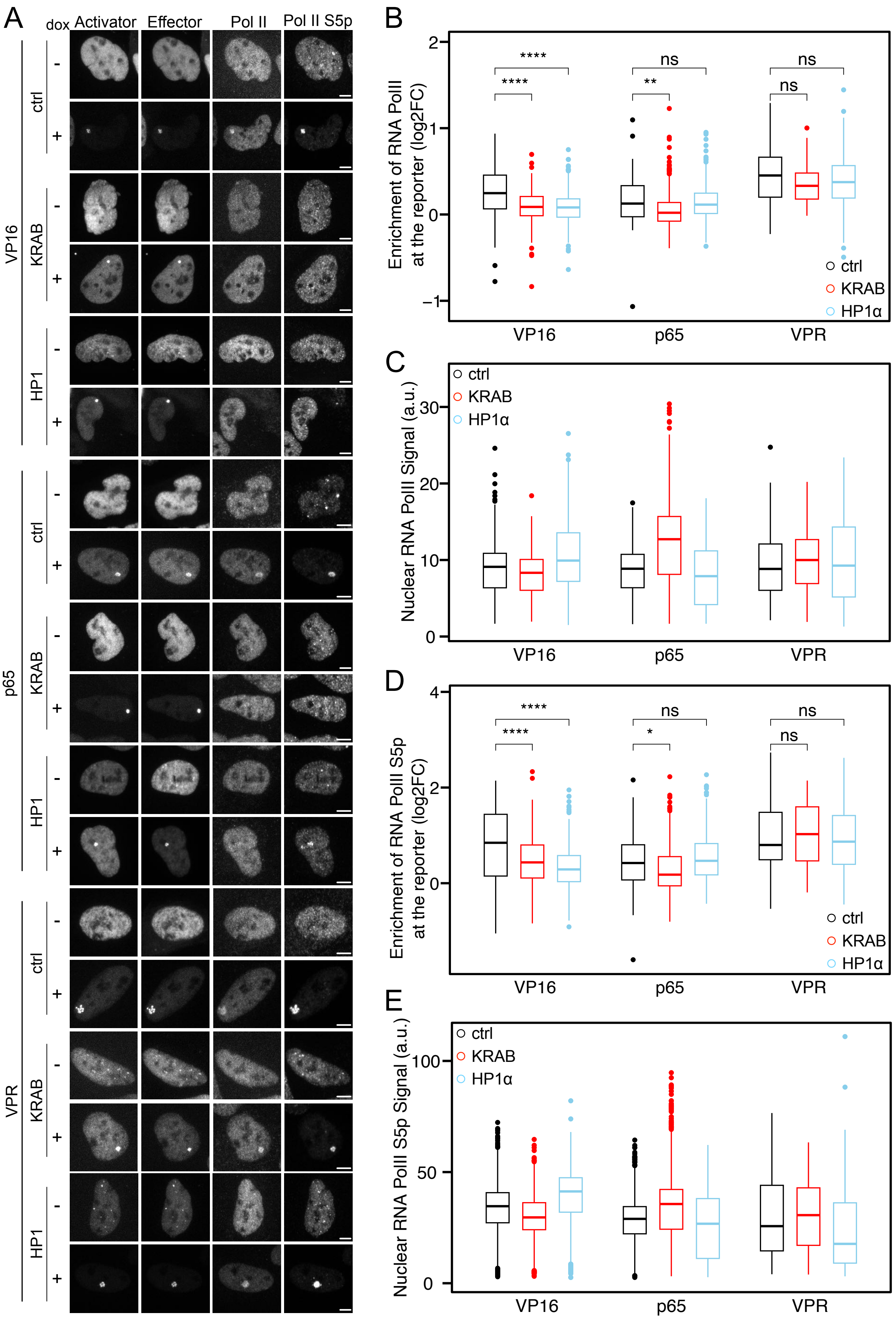
A
dox
Activator
Effector
Pol II
Pol II S5p
VP16
p65
VPR
ctrl
KRAB
HP1
-
+
B
Enrichment of RNA PolII at the reporter (log2FC)
****
****
ns
**
ns
ns
ctrl
KRAB
HP1α
VP16
p65
VPR
C
Nuclear RNA PolII Signal (a.u.)
ctrl
KRAB
HP1α
VP16
p65
VPR
D
Enrichment of RNA PolII S5p at the reporter (log2FC)
****
****
ns
*
ns
ns
ctrl
KRAB
HP1α
VP16
p65
VPR
E
Nuclear RNA PolII S5p Signal (a.u.)
ctrl
KRAB
HP1α
VP16
p65
VPR

**Figure S2. HP1α-dependent changes of RNAP II abundance and CTD S5p at the reporter.** (**A**) Confocal microscopy images showing co-recruitment of the activator with HP1α or KRAB and its effect on Pol II and Pol II S5p after 24 h post-doxycycline addition. Scale bar, 5 µm. (**B**) Quantification of the log2FC of RNA Pol II signal at the reporter compared to the nucleoplasm outside at 24 h post-doxycycline addition. n = 51-443, per combination of activator and HP1α or KRAB or control across two biological replicates. Wilcoxon test, **p < 0.01, ****p < 0.0001, p > 0.05 not significant (ns). (**C**) Nuclear RNA Pol II levels were quantified for control samples without doxycycline. n = 243-1491 per combination of activator and HP1α, KRAB, or control across two biological replicates. (**D**) Quantification of the log2FC of RNA Pol II S5p signal at the reporter compared to the nucleoplasm outside at 24 h post-doxycycline addition. n = 51-443, per combination of activator and HP1α or KRAB or control across two biological replicates. Wilcoxon test, *p < 0.05, ****p < 0.0001, p > 0.05 not significant (ns). (**E**) Nuclear RNA Pol II S5p levels were quantified for control samples without doxycycline. n = 273-1491, per combination of activator and HP1α, KRAB, or control across two biological replicates.

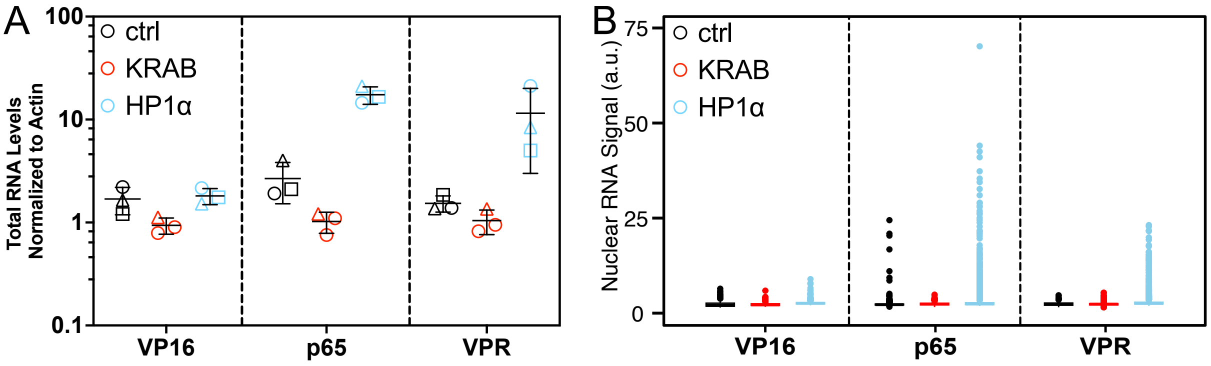


**Figure S3. Uniform transcription levels in the absence of doxycycline across bulk and single-cell readouts** (**A**) Quantification of total RNA levels by RT-qPCR at 24 h for the control with no doxycycline. Mean and SD of fold-change induction normalized to beta-actin mRNA x 1000 (n = 3). (**B**) Nuclear RNA levels were determined at 24 h for the control with no doxycycline, n = 1103-3283, per combination of activator and HP1α or KRAB or control across two biological replicates.

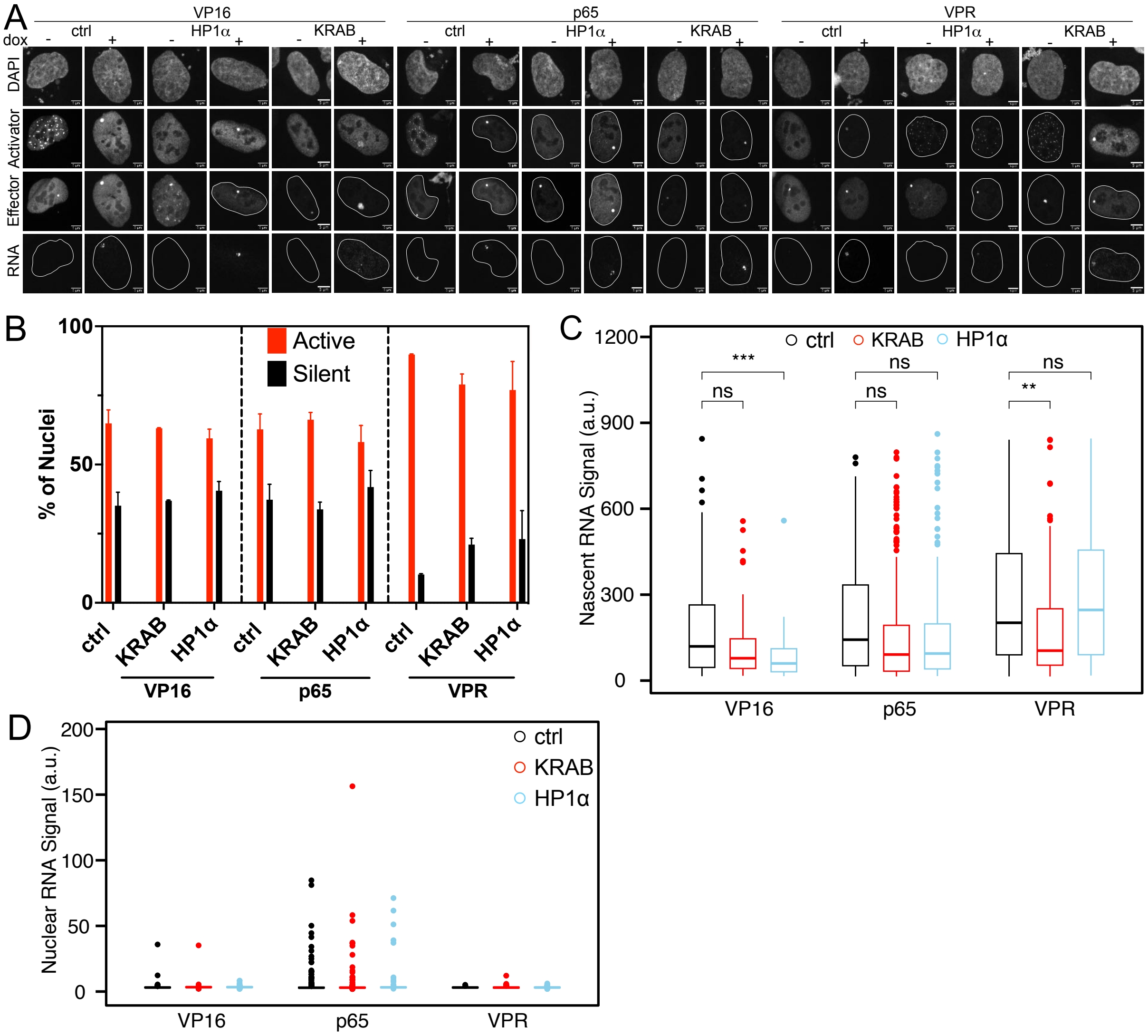


**Figure S4. Dependence on promoter proximity for the repressive effect of HP1α demonstrated at a single-cell level** (**A**) Confocal microscopy images showing light and doxycycline induced recruitment of the activator to the *tet*O repeats for 6 h after 48 h of LacI mediated recruitment of HP1α or KRAB to *lac*O repeats and its effect on transcription, as detected by RNA-FISH. Scale bar, 5 μm. (**B**) Fraction of active or silent nuclei. The classification was based on the presence or absence of transcripts detected by RNA-FISH. Bar: mean, error bars: minimum and maximum of two biological replicates. (**C**) Nascent RNA levels were determined for the nuclei within the active category shown in panel B. n = 59-266, per combination of activator and HP1α , KRAB, or control across two biological replicates. Wilcoxon test, *p < 0.05, ***p < 0.001, p > 0.05 not significant (ns). (**D**) Nuclear RNA levels were determined by RNA-FISH after 48 h of HP1α or KRAB recruitment to *lac*O repeats followed by 6 h of light exposure for the control with no doxycycline, n = 324-1430, per combination of activator and HP1α or KRAB or control across two biological replicates.

p65
repressed

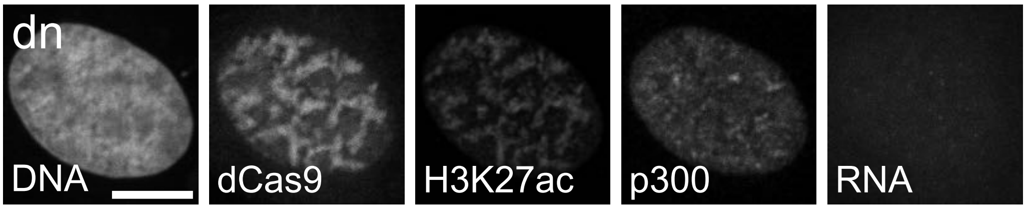


**Figure S5. Example of *Suv39h* dn p65 repressed phenotype.** Roughly 2% of cells represent this phenotype and show decondensed chromocenters but no transcription activation. Scale bar, 10 µm.
